# LoAR-Exo and HiAR-Exo: One-step label-free isolation of extracellular vesicles using inertial microfluidic devices

**DOI:** 10.1101/2025.10.10.681648

**Authors:** Sourav Acharya, Moein Talebian Gevari, Ashwin Anandhapadman, Shreya Karandikar, Haimanti Mukherjee, Fredrik Stridfeldt, Sayani Das, Petra Hååg, Nupur Agarwal, Rolf Lewensohn, Kristina Viktorsson, Sandip Kaledhonkar, Prakriti Tayalia, Apurba Dev, Debjani Paul

## Abstract

Extracellular vesicles (EVs) are lipid membrane-bound nanoscale (20 nm – 1000 nm) objects shed by all cells. As EVs play an important role in intercellular signaling, these have emerged as promising biomarkers for many diseases including cancer and neurodegenerative disorders. A major bottleneck in research into EVs and their smooth translation to clinic as disease biomarkers is a lack of access to affordable and user-friendly technologies to quickly isolate EVs from complex biological and clinical samples with acceptable purity and yield. Here, we report two different designs of inertial microfluidic devices (e.g. LoAR-Exo and HiAR-Exo) that can isolate EVs from different kinds of biological samples in a single label-free step using a spiral microchannel design with different aspect ratios. LoAR-Exo has a height-to-width aspect ratio ∼ 1, while HiAR-Exo has a height-to-width aspect ratio of 2.5, optimized by COMSOL simulations. We tested both designs by separating polystyrene particles of size < 1 μm from a heterogeneous mixture. We then benchmarked the performance of the microfluidic chips against ultracentrifugation and a precipitation kit by isolating EVs from the cell culture-conditioned media (CCM) of MDA-MB-231 cells. We also compared the microchip with SEC by using the CCM from H1975/OR cells. The size distribution of EVs isolated by the microfluidic chip was comparable with ultracentrifugation, precipitation and SEC. In summary, both devices isolated EVs from as little as 1 mL of sample volume using a label-free technique in a continuous manner and without any user intervention. Both microfluidic platforms offer a simple, efficient, and scalable alternative to conventional methods for EV isolation.

## Introduction

Extracellular vesicles (EVs) are nanoscale biological particles released by cells and found in body fluids [Van Niel et al. 2018]. They play a vital role in cellular functions, such as cell to cell communication, tissue maintenance and response to inflammatory and traumatic injuries [Mathieu et al. 2019]. EVs are promising biomarkers for different diseases because they carry important cargo molecules, including mRNA, lipids, and proteins, that convey information about their origin. EVs are highly diverse in size (20 - 1000 nm), composition of surface proteins, and origin [Welsh et al. 2023]. However, studying EVs is challenging due to difficulties in isolating and analyzing individual vesicles, given their small size and the complexity of separation techniques as well as due to the heterogeneity of the EVs when isolated from bodily fluids.

While small EVs (20 nm – 200 nm) have been widely studied, large EVs ranging in size from 200 nm to 1 µm are now also gaining a lot of interest for diagnostic applications. This is because EVs in this size range have a higher number of membrane receptors and contain a larger amount of cargo due to their larger size [Nada Ahmed et al. 2025], which also makes them interesting to study. Most EV-isolation technologies focus on isolating small EVs, often overlooking the significance of larger EVs as a source of disease-linked markers.

A crucial step in isolating EVs from complex biological samples is eliminating larger micron-sized contaminants. Ultracentrifugation (UC), which involves several successive centrifugation steps, has for long been the gold standard EV isolation technique. It has several limitations, such as damage to the protein corona of the EVs and increase in production of multi-lamellar vesicles, long processing times (∼ 6 - 8 h), requirement of large sample volumes (∼ several millilitres) to get a pellet at each centrifugation step, and the loss of the EVs material in each successive centrifugation step [Lobb et al. 2015]. Size exclusion chromatography (SEC) is another widely used method that separates particles based on size by passing them through a bed of porous beads [Boing et al. 2014]. Chromatographic techniques such as SEC, size-exclusion liquid chromatography, and anion exchange chromatography have emerged as effective methods for EV isolation, offering advantages in both purity and yield in particular when working with small volume of bodily fluids [Stam et al., 2021; Koch et al., 2024; Kapoor et al., 2024]. However, other particles of similar sizes as EVs can interfere with separation, leading to longer processing times and requiring further purification steps [Janine Stam et al. 2021]. For example, Bergqvist et al. utilized a combination of superabsorbent polymer-based isolation and SEC to achieve a more refined and efficient extraction of EVs from conditioned media of HEK293 cells [Bergqvist et al. 2025]. Compared to conventional ultracentrifugation, multimodal chromatography has demonstrated improved purification of EV populations from blood plasma [Zimmerman et al., 2024]. Precipitation kits have emerged as another alternative to ultracentrifugation as they can separate EVs in fewer steps yet many of the precipitation kits need an overnight incubation step [Schageman et al. 2013], increasing the total processing time. The polymers used to precipitate EVs in these kits are very difficult to remove, thereby compromising the purity of the isolated EVs [Clos-Sansalvador et al. 2022] and also prohibiting proper downstream characterization of particle size and/or cargo. Importantly, it should be noted that all these methods work with fixed sample volumes and cannot isolate EVs continuously.

Inertial microfluidic technologies, with Reynolds number, Re > 1, offer a promising solution for continuous label-free isolation of sub-micron particles and EVs using only hydrodynamic forces (i.e. lift and Dean drag forces). Researchers have used curved microchannel designs for effective inertial focusing of EVs [Tay et al. 2021, Leong et al. 2024]. However, as both Exo-DFF [Tay et al. 2021] and ExoArc [Leong et al. 2024] use pinch flow fractionation to separate the particles, it requires a flow-rate as high as 400 µL/min that may damage the particles. Additionally, viscoelastic forces combined with inertial forces in complex microchannel designs have been employed to enhance the focusing of particles of EVs sizes [Liu et al. 2017, Shiri et al. 2022, Zhou et al. 2019, Tanriverdi et al. 2025]. In particular, Tanriverdi et al. used elasto-inertial flow in a high-aspect ratio (∼ 6) straight microchannel to focus down to 20 nm particles and EVs (100 nm). These particles applied to demonstrate focussing were isolated using a combination of multiple centrifugation steps, and tangential flow filtration. Sample contamination remains a significant challenge in such systems, primarily due to the difficulty of removing residual viscoelastic material following the isolation [Lan et al. 2025]. It should be noted that several research groups have demonstrated separation of non-biological sub-micron particles using inertial microfluidics alone or in combination with other hydrodynamic forces, but have not extended their technique to separation of biological particles in the same size range (such as EVs) [Mutlu et al., 2018, Wang et al., 2017, Kim et al., 2012, Liu et al., 2016]. Cruz et al reported a High-Aspect Ratio Curved (HARC) microchannel design and simulated deterministic focussing of particles ranging in size from 0.7 µm to 1 µm [Cruz et al, Jl. Micromech. Microengg. 2021]. They then developed the theoretical framework for focussing of particles of different sizes in this HARC design [Cruz et al, Sci. Rep. 2021]. Finally, they experimentally demonstrated separation of polystyrene particles ranging in size from 790 nm to 1 µm, at 80 nm resolution [Cruz et al, Sci. Rep. 2021] in HARC. However, they did not extend the HARC concept to separation of EV-like particles, which requires handling of complex biological samples having particles of different sizes ranging from ∼ 10 nm to tens of micrometers.

Here we report two variations of a spiral microfluidic device designed to continuously separate all particles <1000 nm in size, including EVs, from complex biological samples in a *single* label-free step (**Figure 1a**) without any pre-incubation or sample processing. Both designs have a single spiral microchannel with an optimized height-to-width (h/w) ratio that preferentially pushes all particles larger than 1 µm towards the inner wall of the channel (**Figures 1b, 1c** and **1d**). Particles smaller than 1 µm are collected from a designated sample outlet abutting the outer wall of the channel. We report two different spiral device designs: (a) LoAR-Exo with an aspect ratio (AR) of ∼ 1, and (b) HiAr-Exo with an aspect ratio of 2.5. Using LoAR-Exo, we isolated nanoparticles (with size < 1 µm) from a heterogeneous mixture of polystyrene beads ranging in size from 100 nm to 1 µm. Next, we isolated EVs from the cell culture-conditioned media (CCM) of breast cancer (BC) (MDA-MB-231) and non-small cell lung cancer (NSCLC) (H1975/OR) cell lines [McGowan M et al. 2017] using the same design. As seen from nanoparticle tracking analyses (NTA) data, the size distribution of the EVs obtained from the H1975/OR CCM using LoAR-Exo was comparable to that obtained by the standard size-exclusion chromatography (SEC). Using a fluorescence-based single-EV detection technology, we found a similar abundance of different tetraspanins in EVs isolated either by our device or SEC. We achieved a 6.7-fold higher yield (1.6 × 10^10^ particles) in LoAR-Exo compared to ultracentrifugation (2.38 × 10^9^ particles) when using CCM of MDA-MB-231 BC, without any damage to the EV structure as seen in the cryo-TEM images. In summary, both our designs can isolate EVs from a wide range of biological samples due to their ability to perform continuous operation.

**Figure 1.**
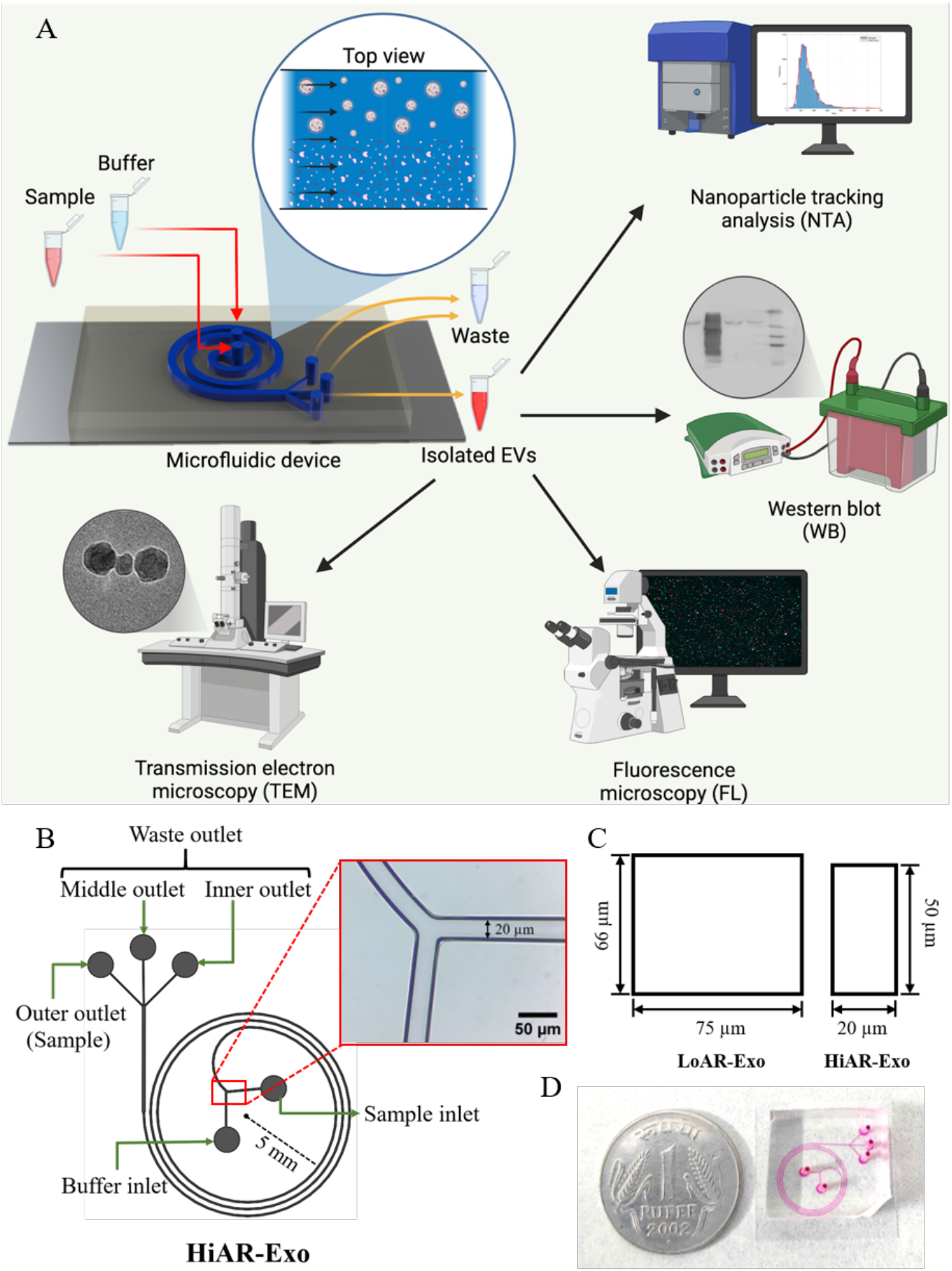
The design of the microfluidic device. (A) A schematic representation showing the process flow of the platform. (B) Schematic diagram of LoAR-Exo and HiAR-Exo device designs. The inset shows a photo of HiAR-Exo. (C) Schematic diagram showing the cross-sectional dimensions of LoAR-Exo and HiAR-Exo. (D) Photo of LoAR-Exo filled with a dye. A 22 mm-dia one-rupee Indian coin is placed next to it to show the channel footprint.

## Materials and Methods

### Design of the microfluidic device

Both LoAR-Exo and HiAR-Exo designs (**Figure 1b** and **1c**) have a spiral channel with two inlets and three outlets. One is the ‘sample inlet’ and the other is the ‘buffer inlet’. The buffer inlet ensures that particles inside the sample are confined near the inner wall of the channel by a sheath flow when they enter the device, which helps to reduce the time required for particle focusing. The channel is finally divided into three outlets of equal width. The provision of a middle outlet ensures that it can either be designated as a sample outlet or as a waste outlet, depending on the size cut-off of the desired particles. In the current study, the inner and the middle outlets together constitute the ‘waste outlet’ for liquid material including particles larger than 1 µm. The outer outlet (or the ‘sample outlet’) is the one from which particles like EVs are collected. LoAR-Exo and HiAR-EXo have different cross-sections and aspect ratios, while keeping the channel length and the radius of curvature the same in both designs. In LoAR-Exo, the aspect ratio of the spiral channel is ∼ 1, while HiAR-Exo has a high (2.5) aspect ratio with a channel width of 20 µm and an optimised height of 50 µm. The average radius of curvature (R) of both spiral microchannels is 5 mm, with 2.75 number of loops. While we largely followed the theory of high-aspect ratio inertial microfluidic devices reported by Cruz et al [Cruz et al, Jl Micromech. Microengg. 2021] while designing HiAR-Exo, HiAR-Exo has three major modifications. First, our design incorporates an additional buffer inlet that facilitates particle constriction near the inner wall. Second, it features a ten-fold larger radius of curvature, which simplifies fabrication. Third, the increased channel width enables efficient focusing of particles in sizes of a few microns.

### COMSOL simulation of fluid flow and particle focussing

We optimized the device dimensions and the flow rate using COMSOL multiphysics software (version 5.4). The density and the viscosity of the solution (water) are assumed to be 1000 kg/m^3^ and 1 cP respectively. We used physics-controlled tetrahedral mesh of size 1.56 µm in the channel. COMSOL used a Newtonian solver with an initial damping factor of 0.001. The solution was considered to be converged for a residual factor of 0.01 or a maximum iteration of 200. We obtained velocity profiles and streamlines at different flow rates and for different geometrical parameters (i.e aspect ratio and radius of curvature). Once the geometry and flow rate were finalised for each design, we simulated the separation of a heterogeneous suspension containing 200 nm, 500 nm, 800 nm, 1 µm, and 3 µm polystyrene particles (10^6^ particles/mL) using the particle tracing module of COMSOL. The density of the polystyrene particles was taken as 1050 kg/m^3^ [Mueller et al. 2020]. We used a time-dependent solver with a time step of 0.001 sec and ran the simulations for 5 sec and 1 sec for LoAR-Exo and HiAR-Exo respectively to obtain the final particle positions across the channel cross-section just before the outlets. We also separately simulated focusing of 1 µm and 200 nm particles in HiAR-Exo for concentrations ranging from 10^4^ particles/mL to 10^7^ particles/mL to see if the increase in concentration affects the focussing.

### Fabrication of the microfluidic devices

The mask design was made using SolidWorks 2021 and the chrome mask was prepared using a Microwriter ML 3 (Durham Magneto Optics, UK) at IIT Bombay’s Nanofabrication Facility (IITBNF). SU-8 lithography was carried out in the cleanroom of IITBNF. SU-8 (MicroChem Corporation, USA) was spin coated (WS-400 BZ from Laurell Technologies, USA) on a 2-inch clean silicon wafer at 1000 rpm for 15 seconds, followed by 3500 rpm for 35 seconds. The wafer was then prebaked on a hot plate at 65°C for 5 min, followed by 95°C for 15 min. The photoresist was then exposed to UV in a mask aligner (EVG 620 from EV Group, Austria). Post-exposure bake was done at 65°C for 1 min and 95°C for 5 min, followed by development for 20 min. The microfluidic devices were fabricated using the elastomer Polydimethylsiloxane (PDMS) (Sylgard 184, Dow Corning, India). The PDMS base was mixed with the curing agent in a 10:1 ratio while casting on the SU8-patterned substrate, and cured it at 65°C for 45 min. The inlets and outlets were punched using a 1.5 mm biopsy punch (Accu-Sharp, Med Morphosis LLP, India). The PDMS devices were then bonded to a clean coverslip (Blue Star, India) using oxygen plasma (PDC 32G from Harrick Plasma, USA) for 90 sec. A final curing step at 65°C was carried out for 40 min to strengthen the PDMS-glass bond.

### Protocol for separation of polystyrene particles

We performed particle separation experiments in parallel at IIT Bombay and Uppsala University. At IIT Bombay, a suspension of polystyrene beads (Sigma-Aldrich, USA) of sizes 1 μm (10^7^ particles/mL), and 500 nm (10^8^ particles/mL) were prepared in filtered distilled water to separate nanoparticles in the HiAR-Exo design. For experiments performed in Uppsala University, using the LoAR-Exo design, the polystyrene beads of sizes 1 μm and 100 nm (both at 10^7^ particles/mL concentration) were suspended in filtered distilled water. 1 mL of the heterogeneous particle suspension flowed through the sample inlet at 50 μL/min (Model 78-8111C, Cole Palmer, India), while filtered distilled water was passed through the buffer inlet at 100 μL/min (GT 1001, Genietouch, USA). Particles were collected from all waste and sample outlets in Eppendorf tubes.

### Cell culture and collection of cell culture-conditioned media for EV isolation

The MDA-MB-231 cells were cultured in DMEM (Sigma Aldrich, USA), supplemented with 10% fetal bovine serum (FBS) (Sigma Aldrich, USA) and 1% antibiotic (10,000 U Penicillin and 10 mg Streptomycin per ml in 0.9% normal saline). A non-small cell lung cancer (NSCLC) cell line, H1975/OR developed to be resistant to epidermal growth factor receptor (EGFR) tyrosine kinase osimertinib [McGowan M et al. 2017] was a kind gift from Professor Odd Terje Brustugun, Vestre Viken Hospital Trust, Drammen and Institute for Cancer Research, Oslo University Hospital, Oslo Norway. The cells were maintained in RPMI-1640 medium supplemented by fetal bovine serum (FBS, 10%) and 2 mM L-glutamine (Gibco, Life Technologies, Stockholm, Sweden). For the experiment the cells were grown in a medium containing exosome depleted FBS (A2720801, Thermo Fisher Scientific).

The decanted media was collected after 48 h of culture when the cell count reaches ∼ 10^5^ cells/mL. This cell culture-conditioned media (CCM) had extracellular vesicles, growth factors, cytokines and metabolites secreted by the cells. The collected CCM was kept on ice and used on the same day.

### Protocol for EV isolation from CCM using LoAR-Exo and HiAR-Exo

Two syringes (BD Scientific, India) were filled with CCM and 1X filtered phosphate buffered saline (PBS) (Gibco, India) respectively, and these were connected to the inlets. CCM and PBS were passed at 50 μL/min and 100 μL/min respectively. The sample and the waste were both continuously collected in eppendorf tubes kept on ice. All experiments were conducted in triplicates.

### Protocol for EV isolation from CCM using ultracentrifugation

48 mL of CCM was collected after 48 h culture of MDA-MB-231 cells when the cell count reaches ∼ 10^5^ cells/mL. The isolation process involved a series of centrifugation steps at 4°C to systematically eliminate cellular contaminants. An initial centrifugation step at 250g for 10 min was used to remove detached cells. The supernatant was collected and centrifuged at 2,500g for 10 min to remove cell debris. To further purify the sample, the supernatant was centrifuged at 15,000g for 20 min to remove microvesicles and apoptotic bodies. For the final ultracentrifugation step, the remaining supernatant was centrifuged at 150,000g for 150 min using an ultracentrifuge (Optima XPN 100, Beckman Coulter, USA) equipped with 70Ti fixed rotors. Following ultracentrifugation, the supernatant was discarded, and the resulting EV pellet was resuspended in 200 μL of 1X PBS.

### Protocol for EV isolation from CCM using a precipitation kit

1 ml of CCM was collected after 48 h culture of MDA-MB-231 cells. A centrifugation step of 2000g was performed on the sample for 30 mins to remove cellular components. 0.5 mL of precipitation reagent (Total Exosome Isolation Reagent, Thermofisher, USA) was mixed with the sample and incubated overnight at 4°C. The sample was then centrifuged at 10,000g for 1 h at 4°C. The supernatant was discarded and the pellet was thereafter resuspended in 1 mL 1X PBS.

### Protocol for EV isolation from CCM using size exclusion chromatography

EVs were isolated from 50 mL of CCM of H1975/OR cells with two steps of centrifugation in a Rotina R38 centrifuge (Hettich, Germany) to clear out cell debris. Initial centrifugation was at 200g for 5 min, followed by centrifugation of supernatant at 720g for 10 min. The CCM was concentrated to ∼ 500 µL using Amicon Ultra-15 Centrifugal Filter Unit with a MWCO of 3 kDa (#UFC900324, Merck Chemicals and Life Science AB, Solna, Sweden). EVs were isolated by SEC on qEV original gen2 columns (Izon Science, Oxford, UK) as previously described [Stiller et al, Cancer, 2021]. Briefly, the samples were added to the column and were eluted in 500 µL fractions by 0.22 µm filtered PBS. Fractions 6-10 were pooled and concentrated to about 500 µL using an Amicon® Ultra-4 Centrifugal Filter Unit (#UFC800324, Merck Chemicals and Life Science AB).

### Measurement of concentration of polystyrene particles and EVs by nanoparticle tracking analysis

At IIT Bombay, the concentration of all polystyrene particles collected from the sample outlet in HiAR-Exo were obtained by using nanoparticle tracking analysis (NTA) on a NanoSight NS300 (Malvern Panalytical, UK) system. NTA measurements were performed using a camera level of 10 and a resolution value of 81. At Uppsala University, the characterization of both 1 μm and 100 nm particles in LoAR-Exo were measured using a Zetaview (Particle Matrix, Germany) system as it has an upper size cut-off of 10 μm. The camera settings were optimized for different particle sizes as presented in the **Supporting Information**. The concentrations of the EVs were measured by NTA using both a NanoSight system and a ZetaView system.The CCM from MDA-MB-231 cells was analyzed on the NanoSight without dilution while EVs isolated from the H1975/OR cells were diluted 1:1000 in sterile, filtered 1× PBS for the ZetaView measurements. In Nanosight measurements, 1 mL of sample was loaded into the channel and the movement of particles was recorded for 1 min. In Zetaview measurements, for each run, 100 µL of sample was loaded and recorded in triplicate across eleven distinct positions. All measurements were performed at 24–25 °C.

### Western blot analyses of EVs from CCM

Western blot analysis was performed on samples collected from the sample outlet of the HiAR-Exo device. For CCM samples from MDA-MB-231 cells, cell lysate was used as a positive control. Prior to western blotting, the total protein concentration in the sample collected from the sample outlet was quantified by Bradford assay (at 595 nm) using bovine serum albumin (BSA) as a standard. Based on the Bradford assay results, a minimum of 35 μg of total protein from the sample was used for western blot. The samples were mixed with loading dye under reducing conditions and run on 12.5% polyacrylamide gels with Tris-glycine running buffer. Proteins were transferred onto PVDF membranes with a transfer buffer containing 20% methanol at 4°C. The membranes were blocked with 5% BSA (MB083, HiMedia, India) in TBS-T (Tris-buffered saline with 0.1 % Tween 20, pH 7.6) followed by overnight incubation at 4°C with primary antibodies against CD63 (Rabbit mAb, Abclonal, USA, cat no. A19023), CD81 (Rabbit mAb, Abclonal, USA cat no. A22528), TSG101 (Rabbit mAb, Abclonal, USA, cat no. A5789), and calnexin (Rabbit mAb, Abclonal, USA, cat no. A4846). All primary antibodies were used at 1:1000 dilution in 0.5% BSA in TBS-T. After incubation, the membrane was washed with TBS-T and a secondary antibody (HRP Goat Anti-Rabbit IgG, Abclonal, USA, cat no. AS014, at 1:10,000 dilution) was added at room temperature for 2 h. After additional washing steps with TBS-T, the blot was developed using the Westar Hypernova kit (Cyanagen, Italy) in both chemiluminescence and colorimetric mode in the Amersham ImageQuant imaging set-up (IQ500, Cytiva, USA).

### Single EV measurement

To characterise the isolated vesicles by the microchips, an LED equipped inverted microscope (Axio Observer 7 from Zeiss) was used to perform single EV analysis. For this purpose, the EVs were captured non-specifically on a silica coverslip equipped with a 4-well reservoir using poly-L-lysine coating. After the EV incubation, the surface was treated by Casein to suppress the non-specific interactions of the fluorophore-tagged antibodies. 4 nM concentration of the detection antibodies were incubated on EVs captured in each well. CD9-VioBlue, CD63-R-PE, and CD81-APC are the three detection antibodies used in this study. The substrate preparation, reagents, measurement, and data analysis followed our previous study [Stridfeldt et al. 2023].

### Data analysis and statistics

All LoAR-Exo and HiAR-Exo experiments were carried out using different CCM from three biological cell replicates. All the experiments with cell-spiked CCM were also performed from three biological replicates. Unless otherwise specified, we report the mean values along with standard error of mean (SEM).

## Results

### Optimization of the device geometry and flow rate by COMSOL simulation

We optimized the geometrical and flow parameters for LoAR-Exo and HiAR-Exo separately. We first simulated the flow in both designs using COMSOL to determine the optimum radius of curvature (R) of the spiral channel needed to generate stable secondary flows. LoAR-Exo had an aspect ratio (AR) of ∼1. As presented in **Section 1A** of the Supporting Information, the minimum flow rate (Q) required to focus particles > 1 µm in LoAR-Exo was ∼ 100 µL/min. Therefore, we simulated the fluid flow through LoAR-Exo at flow rates of 100 and 150 µL/min. **Figure 2A** shows the Dean vortices that were formed at both flow rates for R = 5 mm. For the HiAR-Exo design, we used the theory of particle focussing in high-aspect ratio inertial microfluidic devices, as shown in **Section 1B** of the Supporting Information. We explored flow-rates of 100 µL/min, 300 µL/min and 500 µL/min for R = 5 mm. We chose R = 5 mm because stable rectangular counter-rotating vortices were created only at Q = 500 µL/min and not for lower flow-rates when R = 10 mm (**Section 3** and **Figure S1** in Supporting Information). **Figure 2B** shows that at R = 5 mm, stable rectangular counter rotating vortices were formed at all flow-rates from 100 µL/min to 500 µL/min. We decided to use R = 5 mm and Q = 150 µL/min (by further imposing a constraint of Q_sample_ = 50 µL/min and Q_buffer_ = 100 µL/min) for further studies.

**Figure 2.**
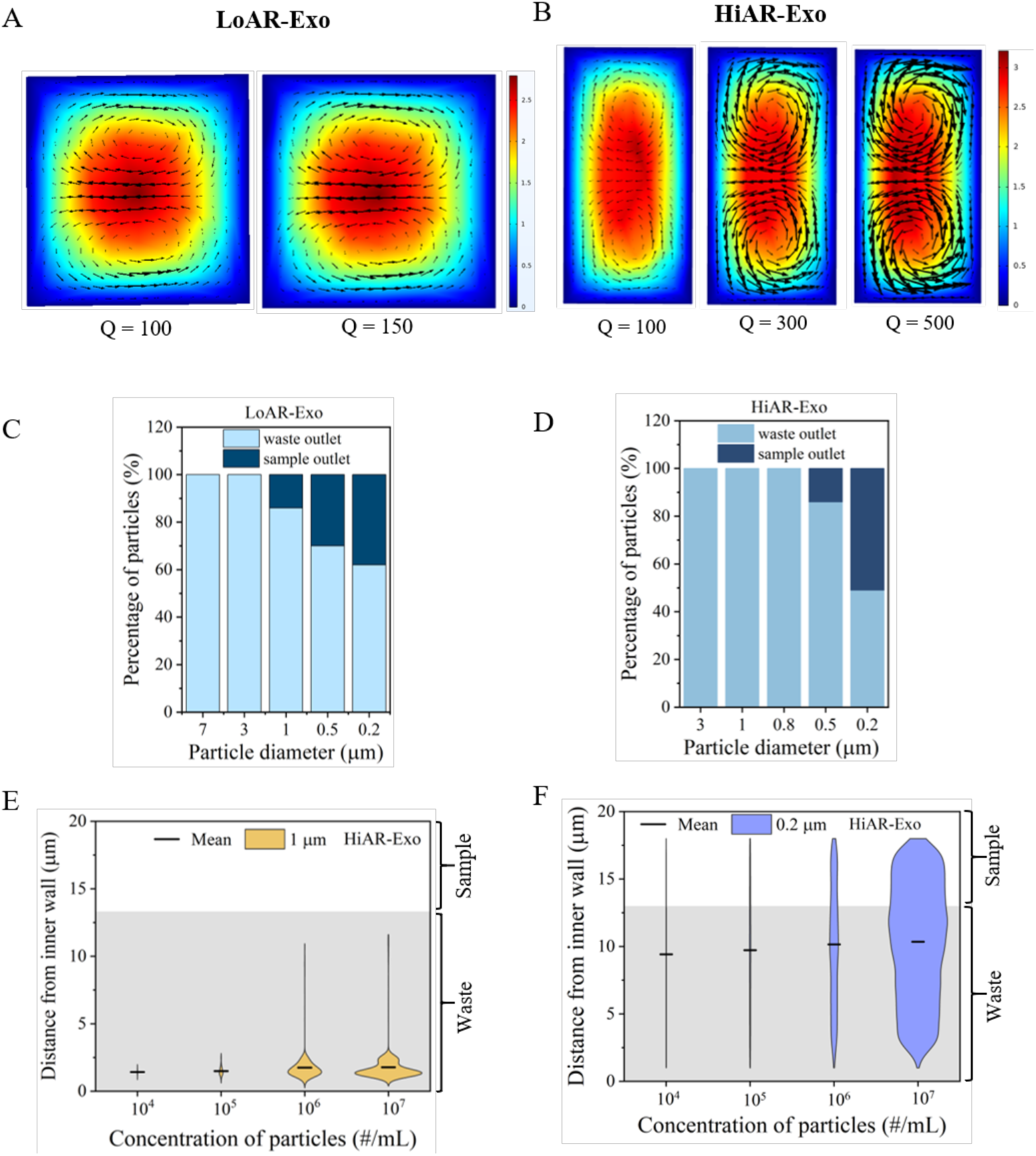
Optimization of geometrical and flow parameters of LoAR-Exo and HiAR-Exo by COMSOL simulation. (A) Cross sectional view of LoAR-Exo showing counter-rotating vortices at R = 5 mm for flow rates of 100 and 150 µL/min. (B) Cross sectional view of HiAR-Exo showing counter-rotating vortices for flow rates of 100 µL/min, 300 µL/min and 500 µL/min. (C)The percentage of the total particles collected in waste and sample outlets of LoAR-Exo. (D) Plot showing percentage of the total particles collected in waste and sample outlets of HiAR-Exo for aspect ratios of 2.5. (E) and (F) show the extent of the focusing of 1 µm and 200 nm particles of different concentrations in HiAR-Exo.

We next used a particle tracing module to analyse the focusing efficiency of LoAR-Exo and HiAR-Exo. We used a sample flow rate of 50 µL/min and a buffer flow rate of 100 µL/min for both devices. In LoAR-Exo, we introduced a heterogeneous sample containing polystyrene particles of sizes 0.2 µm, 0.5 µm, 1 µm, 3 µm and 7 µm and determined the positions of particles of different sizes along the cross-section. We calculated the percentage of particles of specific sizes collected in the sample (dark blue) and the waste (light blue) outlets. As shown in **Figure 2C**, all particles of size 7 µm and 3 µm left the device through the waste outlets. In contrast, ∼12% of 1 µm particles, ∼30% of 0.5 µm and ∼40% of 0.2 µm particles were retained in the sample outlet.

In HiAR-Exo, we analysed the focusing efficiency at different aspect ratios (AR) at R = 5 mm and Q = 150 µL/min. To do this, we varied the aspect ratio (AR) by keeping a fixed value of w = 20 µm, while changing h as 40 µm, 50 µm, 60 µm, 80 µm, and 100 µm respectively. Our aim was to determine at which AR one can focus larger (≥ 1 µm) particles towards the inner wall of the microchannel. We next introduced polystyrene particles of sizes 0.2 µm, 0.5 µm, 0.8 µm, 1 µm, and 3 µm into HiAR-Exo and determined their positions. The distance of individual particles of different sizes from the inner wall and the bottom surface of the channel are shown as violin plots in **Figure S2** and **S3** respectively in the Supporting Information. As seen in **Figure S2**, none of the particles were tightly focussed for AR = 3, 4 and 5. In contrast, for AR = 2 and 2.5, all particles of size ≥ 1 µm were tightly focused near the inner wall. We then calculated the percentage of particles collected in sample (dark blue) and waste (light blue) outlets respectively for AR = 2.5. As shown in **Figure 2D**, all particles of size 3 µm, 1 µm, and µm leave the device through the waste outlets. In contrast, ∼16% of 0.5 µm particles and ∼52% of 0.2 µm particles were collected in the sample outlet. Given these findings, we decided to continue with AR = 2.5 since at this aspect ratio HiAR-Exo should be able to collect more particles i.e. EVs of all relevant sizes (< 1 µm) in its sample outlet.

We also carried out further simulations of HiAR-Exo by varying the concentration of 200 nm and 1 µm particles from 10^4^ to 10^7^ particles/mL to see the effect of particle concentration on the focusing efficiency of our device. As shown in **Figure 2E** and **2F**, all the 1 µm particles were collected in the waste outlets, while ∼ 35% - 40% of the 200 nm particles were collected via the sample outlet for all concentrations explored in our simulations.

### Experiments with polystyrene particles in LoAR-Exo and HiAR-Exo

We next conducted experiments with polystyrene particles in both LoAR-Exo and HiAR-Exo respectively (**Figure 3**). Experiments using LoAR-Exo were conducted with 100 nm and 1 μm particles (**Figure 3A**), while experiments with HiAR-Exo were done using 500 nm and 1 μm polystyrene particles (**Figure 3B**). **Figure 3A** shows the NTA results from LoAR-Exo with a peak at ∼ 128 nm. **Figure 3B** shows the particle size distribution of the particles in the sample outlet of HiAR-Exo with a major peak at 415 nm. In both cases, the peaks of the NTA distribution (black curves) confirm that no large (1 μm) particles were present in the sample outlet of HiAR-Exo. The green curve in **Figure 3A** shows the size distribution of the particles in the waste outlet, with a peak at 961 nm.

**Figure 3.**
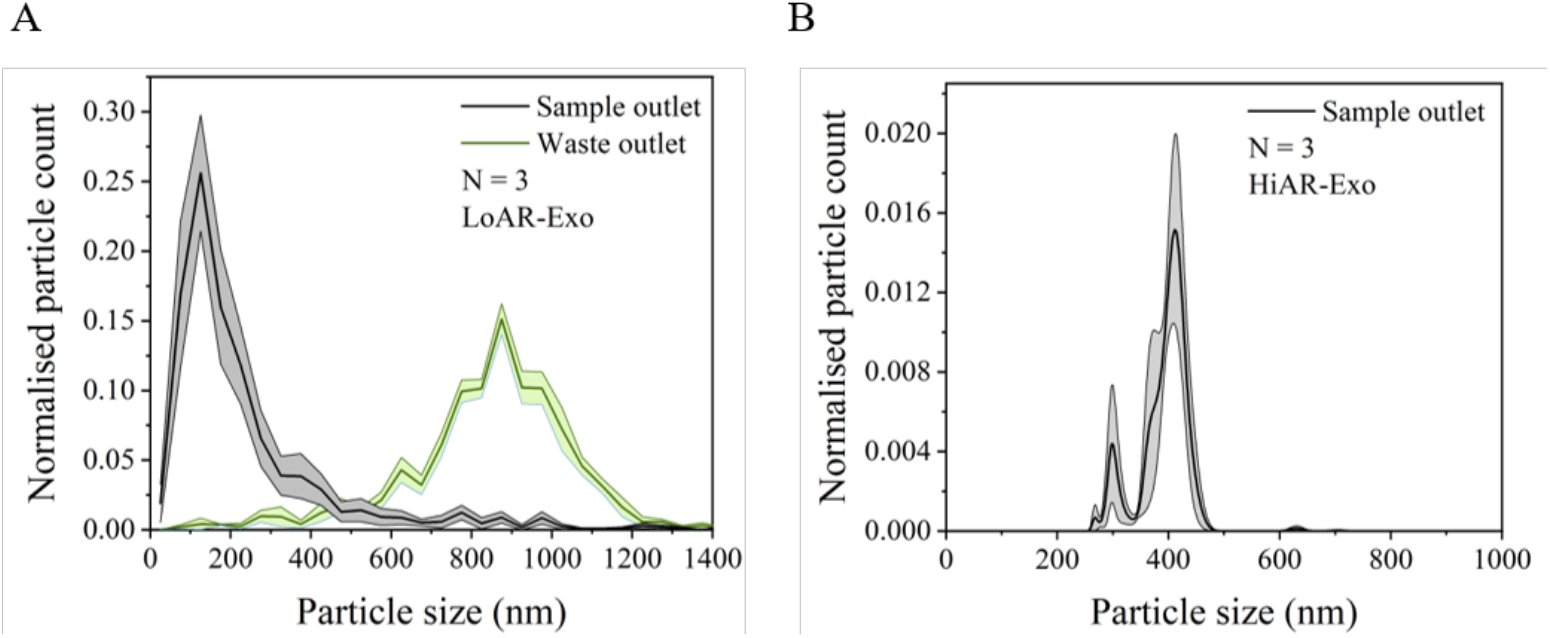
Experiments with polystyrene beads. (A) Size distribution of beads from both sample and waste outlets of LoAR-Exo in experiments (N = 3) done with a mixture of 100 nm and 1 µm beads. (B) Nanoparticle tracking analysis (NTA) plot of HiAR-Exo showing the size distribution of beads (N = 3) in the sample outlet.

### Benchmarking of LoAR-Exo against ultracentrifugation and precipitation methods for EV isolation from cell-conditioned culture media

After testing LoAR-Exo with polystyrene particles, we next wanted to benchmark it against ultracentrifugation (UC) and a commercial precipitation kit as isolation methods with cell culture-conditioned media (CCM) of the breast cancer cell line MDA-MB-231 as a source of EVs. **Figures 4A** and **4B** compare the NTA results from LoAR-Exo against UC and precipitation kit respectively. For these comparisons, the different isolation procedures (microfluidic using LoAR-Exo, UC and precipitation respectively) were done on the same day for each biological sample. As seen, the NTA peak for LoAR-Exo ranges in size from 30 nm to 486 nm with a peak at 119 nm, whereas NTA from UC had a size range from 42 nm to 590 nm with a peak at 123 nm. Thus, both UC and LoAR-Exo showed similar performance in terms of the size of the particles they could isolate, with UC taking ∼ 7 h for the entire process in contrast to just 15 min required for the LoAR-Exo protocol. The total number of EVs as given by the NTA assessment, isolated by LoAR-Exo and UC were (7.7 ± 0.4) x 10^8^ (mean ± SEM, N = 3) and (2.38 ± 0.05) x 10^9^ respectively. It should be noted that UC needed a minimum volume of 22.5 mL of MDA-MB-231 CCM to give a pellet, whereas all LoAR-Exo experiments were performed with only 1 mL of CCM. If the ‘yield’ (i.e. area under the curve) of each technique were normalized by considering equal volumes, the yield of LoAR-Exo was (1.6 ± 0.65) x 10^10^, while the yield of UC was (2.38 ± 0.05) x 10^9^. Therefore, LoAR-Exo had a better yield than ultracentrifugation when equal CCM volumes were considered.

**Figure 4.**
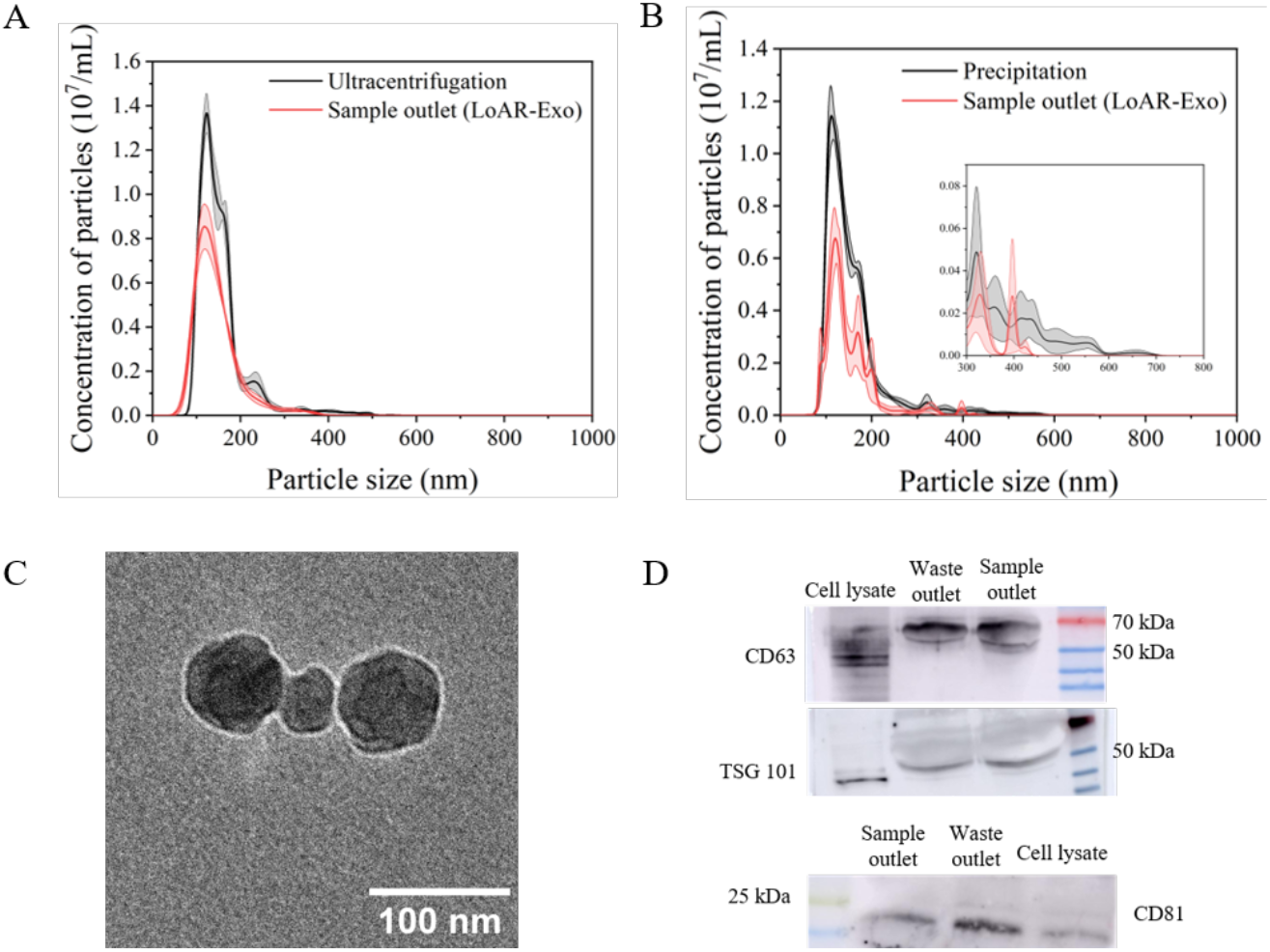
Experiments with cell culture-conditioned media (CCM) sample from the MDA-MB-231 cell line. (A) and (B) compare the EV concentration obtained from LoAR-Exo with ultracentrifugation and precipitation respectively. The inset in (B) shows the particle concentrations from 300 nm to 800 nm. (C) Cryo-TEM image of EVs obtained from LoAR-Exo. (D) Western blot of the EV sample isolated by LoAR-Exo.

**Figure 4B** compares NTA results from LoAR-Exo and the precipitation kit using equal volumes of the CCM samples. The size of EVs isolated by the precipitation kit ranges from 48 nm to 720 nm, whereas EVs isolated by LoAR-Exo on the same day range in size from 42 nm to 470 nm. The yields of EVs isolated by precipitation and LoAR-Exo were (8.7 ± 0.25) x 10^8^ and (4.5 ± 0.18) x 10^8^ respectively. It should be noted that the EV pellet from precipitation also includes the co-precipitated polymer contaminants, as they were difficult to remove prior to downstream applications. Cryo-TEM imaging (**Figure 4C)** of EVs from LoAR-Exo confirms the characteristic cup-shaped morphology. For western blot, two proteins of the category I (CD63 and CD81) and one of the category II (TSG101) markers from the list provided in the MISEV2023 guidelines (Welsh et al. 2024) were selected. **Figure 4D** shows the western blot results for both sample and waste outlets, with cell lysate from MDA-MB-231 applied as positive control. **Figure S4** in the Supporting Information shows the corresponding full images of the western blots. Overall, the experiments with CCM confirm that EVs were present in both sample and waste outlets of LoAR-Exo, while all larger particles were removed from the sample through the waste outlet.

### Benchmarking of LoAR-Exo against size exclusion chromatography-based isolation using a single EV detection technique for evaluation

To further assess the performance of the LoAR-Exo microchip for EV isolation, we used CCM from H1975/OR, in the microchip. In parallel, we applied the same CCM to isolate EVs using size exclusion chromatography (SEC) to benchmark the performance of LoAR-Exo against it. **Figure 5A** shows the normalized size distribution of the EVs isolated by the microchip and SEC. As can be seen, the size distribution of EVs from the microchip and SEC isolation were similar.

**Figure 5.**
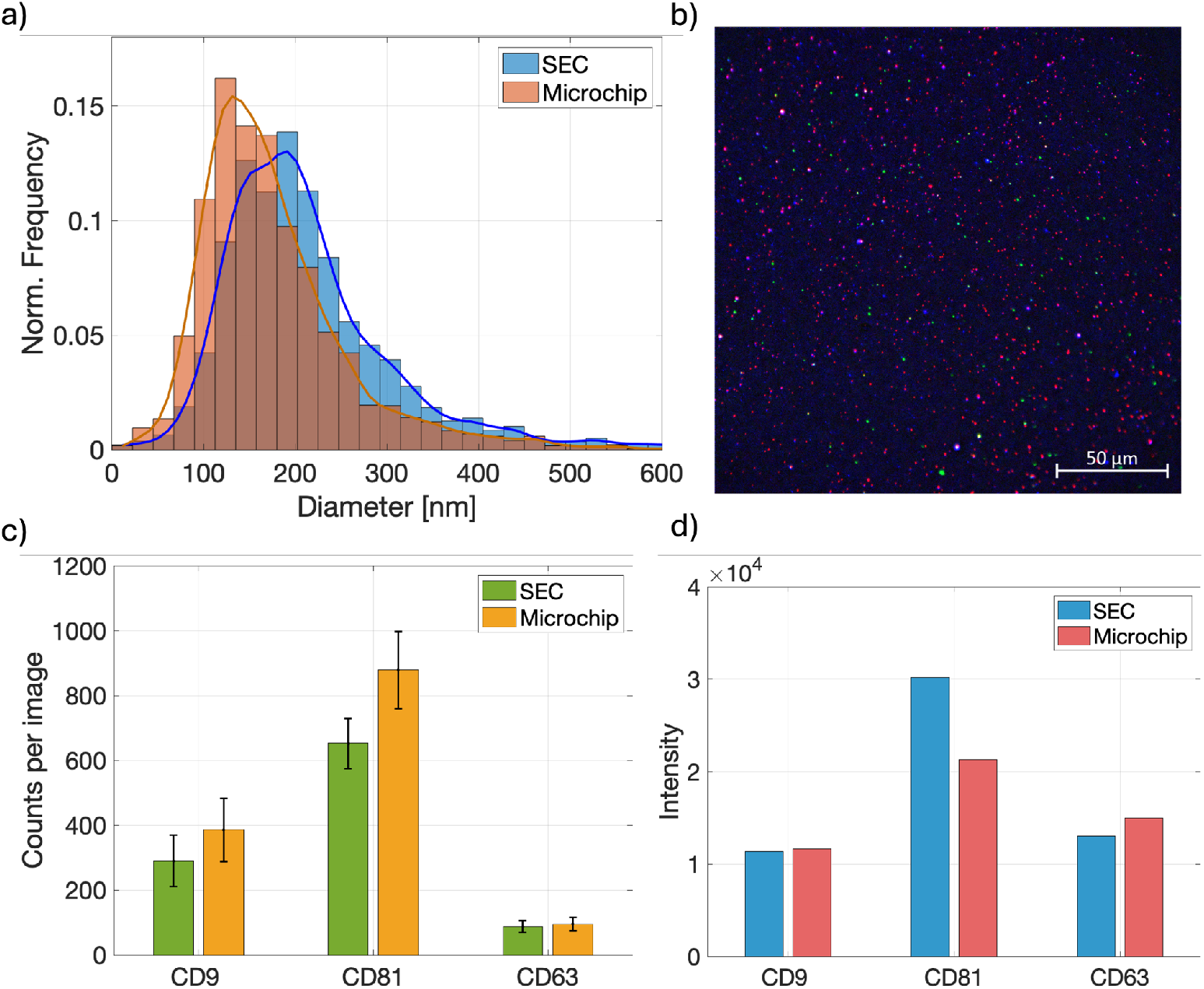
EV isolation from H1975/OR cell culture-conditioned media by LoAR-Exo and size exclusion chromatography. (A) Normalized size distribution of the EVs isolated by either SEC or microchip. (B) Representative fluorescence image of the isolated vesicles by LoAR-Exo. (C) Average EV count per microscopic image after using either of the two isolation techniques. (D) Mean average fluorescence intensity, corresponding to the expression level of the tetraspanins, of the EVs detected by the fluorescence microscope.

Furthermore, we profiled and compared the transmembrane proteins of the isolated EVs using a Zeiss inverted epi-fluorescence microscope under LED excitation. The substrate used to capture EVs was prepared as we previously described [Stridefeldt et al. 2023]. EVs isolated by both LoAR-Exo and SEC were separately analyzed following the same procedure. **Figure 5B** shows a representative fluorescence image of the EVs. In this figure, CD81, CD9, and CD63 tetraspanins are depicted in red, blue, and green artificial colors, respectively. The average count of the detected vesicles in 10 images is shown in **Figure 5C**. Finally, the mean average intensity of the detected spots by the microscope is shown in **Figure 5D** for both isolation techniques. The fluorescence intensity indicates the expression level of the tetraspanins on individual EVs. As can be seen, the detected EVs by both methods showed similar trends in the expression levels of the tetraspanins.

### EVs can be isolated from a cell suspension using LoAR-Exo

After validating the performance of LoAR-Exo on CCM and benchmarking its performance against ultracentrifugation, precipitation and size-exclusion chromatography, we next examined the ability of the microchip to isolate EVs from a cell suspension which have a more complex composition. For this purpose, the CCM was spiked with 3 × 10^5^ MDA MB-231 cells/mL which subsequently was passed through the microchip to isolate EVs from the cell suspension. The number of cells in both sample and waste outlets was counted to obtain the cell concentration in each outlet. As shown in **Figure 6A**, there were no cells present in the sample outlet in any of the experiments. **Figure 6B** shows the NTA results from the sample fraction collected from the sample outlet. Their sizes ranged from 20 nm to 430 nm with an average of 122 nm. This size distribution falls well within the size range of EVs [Welsh et al. 2024]. Western blot analysis was performed using CD63 as an EV marker while calnexin, a protein found in the endoplasmic reticulum, was used to indicate the presence of cell contamination. We also used a cell lysate of MDA-MB-231 as a positive control. As expected, bands for CD63 were present in both sample and waste outlets. Importantly, while distinct bands for calnexin were observed in both the cell lysate and the waste outlet, which contained the spiked cells, no calnexin bands were detected in the sample outlet indicating that these were free of cell contamination yet having CD63 positive EVs. **Figure S5** in the Supporting Information shows the corresponding full images of the western blots. All in all, this experiment illustrates that LoAR-Exo can isolate EVs from a cell suspension in a single step without major cellular contamination.

**Figure 6.**
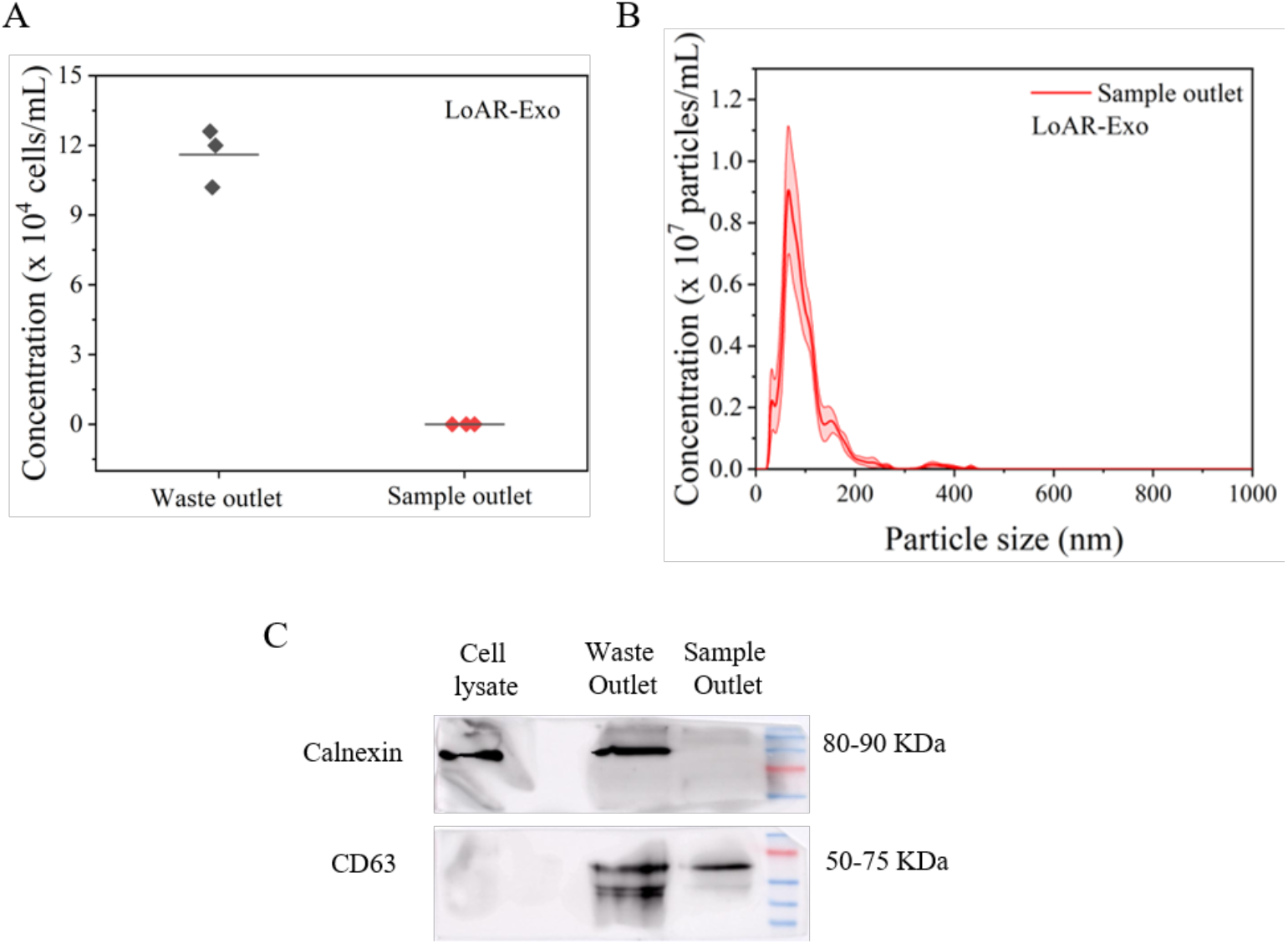
Isolation of EVs from a cell suspension using LoAR-Exo. (A) Plot showing cell concentration in the sample fractions collected from each outlet. Data from three independent experiments are shown, with the dashed line indicating the mean value. (B) NTA plot showing the size distribution of particles collected from the sample outlet. (C) Western blot analysis of the samples collected from sample and waste outlets. A total extract of the MDA-MB-231 cell was used as a positive control.

## Discussion

We report two inertial microfluidic devices (LoAR-Exo with AR ∼ 1 and HiAR-Exo with AR∼ 2.5) to isolate EVs of all sizes from different biological samples in a single label-free step using a spiral microchannel design with different aspect ratios. EV isolation in earlier inertial microchip designs (Exo-DFF [Tay et al. 2021] and ExoArc [Leong et al. 2024]) required strong pinching of the sample flow, which is not the case for LoAR-Exo or HiAR-Exo. Moreover, successful EV isolation in the earlier designs required fabricating the exact channel length at which smaller particles would arrive at the inner wall of the microchannel. In contrast, in both LoAR-Exo and HiAR-Exo, all particles of size > 1 µm remain focussed near the inner wall beyond a minimum channel length required to complete one Dean cycle.

We benchmarked the performance of the LoAR-Exo microchip against UC, SEC and a commercial precipitation kit using cell-conditioned media (CCM). The concentration of EVs isolated from CCM by LoAR-Exo was comparable to UC, while using 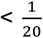 th of the sample volume. Based on our results, we believe that LoAR-Exo is suitable for isolating EVs from samples that have a low abundance (∼ 10^7^ - 10^8^ /mL) of EVs, such as CCM of certain cell lines, saliva, etc [Sjoqvist et al. 2023, Rorhborn et al. 2024, Marques et al. 2024]. Due to the low abundance of EVs in such samples, a large sample volume is initially needed just to get a pellet for example when using UC for isolation. It should be noted that as LoAR-Exo is a flow-based technique, it is equally capable of isolating EVs from a large sample volume by using continuous operation. Furthermore, as UC involves repeated centrifugation steps with increasing g-forces in its purification process, EVs may get damaged [Broad K. et al. 2023]. In contrast, the microchip isolation technique developed in this work minimizes shear damage to EVs. LoAR-Exo gave a size distribution of EVs comparable to UC, SEC and precipitation. Precipitation and LoAR-Exo had almost comparable yields. However, the pellet obtained during precipitation also contained polymer contaminants that were difficult to remove for downstream applications. **Table 1** shows a detailed comparison of LoAR-Exo with these techniques for isolating EVs from CCM. Moreover, as seen in our experiment, the LoAR-Exo design can also be used to isolate EVs directly from cell suspensions further speeding up EV isolation procedures.

**Table 1.** Comparison of LoAR-Exo with other commercial EV isolation techniques. Presented data is based on this work.

| Method | Yield<br>(total<br>particle<br>count<br>in NTA) | Process<br>time (h) | Min.<br>CCM<br>Vol.<br>(mL) | Ease of<br>use<br>(numbe<br>r of<br>steps) | Continuous<br>operation<br>possible? | Additional<br>equipment<br>required? |
| --- | --- | --- | --- | --- | --- | --- |
| Ultra-<br>centrifugation | $10^8$ | 7 | 22.5 | Low<br>(4-5<br>steps) | No | Ultracentrifuge<br>& centrifuge |
| Size<br>exclusion<br>chromatography | $10^{10}$ | 3 | 1 | Low (3<br>steps) | No | Centrifuge &<br>chromatography<br>columns/fraction<br>collector |
| Precipitation kit | $10^9$ | 16<br>(includes<br>overnight<br>incubation) | 1 | Low (3<br>steps) | No | Centrifuge.<br>Specific kit<br>needed for<br>specific sample<br>type. |
| LoAR-Exo (our<br>method) | $10^9$ | 0.25 | 1 | High<br>(one<br>step) | Yes | Two syringe<br>pumps |

On the other hand, HiAR-Exo can directly isolate EVs from cell-free samples, but cannot handle cell suspensions. It can also isolate nanoparticles from a heterogenous mixture of particles. While the HiAR-Exo microchip design is inspired by another high-aspect ratio curved (HARC) microchannel device [Cruz et al. 2021], we made three specific design improvements. To summarize its differences with HARC, the HiAR-Exo design has (a) an additional inlet for buffer, (b) a 10-times higher radius of curvature optimized by simulations, and (c) a wider channel to handle a large range of particle sizes.

Both LoAR-Exo and HiAR-Exo designs preserve the composition of the biological samples containing EVs by only removing the large particles (≥1 µm) while retaining the entire EV population. Currently large EVs (∼ 200 nm - 1 µm, MISEV 2023) are also generating much interest in diagnostic applications as they have a higher number of membrane proteins and can carry more cargo [Nada Ahmed et al. 2025]. In this work we show that our microchip devices also can handle samples in which such large EVs are in focus.

In summary, both LoAR-Exo and HiAR-Exo are promising one-step label-free techniques for continuous EV isolation from a wide range of biological samples with yields comparable to the gold standard techniques. These methods are compatible with other downstream sensing methods. In the future, we propose to develop our platforms to isolate specific sub-populations of EVs.

## Acknowledgements

DP thanks Wadhwani Research Centre for Bioengineering (WRCB) for funds received through their 9th and 12th calls for projects (grants numbers DO/2025-WRCB002-105 and DO/2022-WRCB002-082). This work was also supported by funding from a Blockchain for Impact-BIOME grant (DO/2024-INCO002-003). DP acknowledges the ‘Seed Funding for Collaboration and Partnership Projects’ (RD/0523-IOE00I0-053) from the Institute of Eminence (IOE) cell of IIT Bombay for initiating the collaboration with Uppsala University and Karolinska Institutet. AD acknowledges the VAIBHAV fellowship (RD/0124-INAE040-001) from the Department of Science and Technology (Govt of India). SA acknowledges the WRCB-HiMedia Entrepreneur-in-Residence fellowship (DO/2024-HIME002) for salary support. The co-workers at Karolinska Institutet were supported by grants from Swedish Cancer Society (# 24 3793 Pj to RL) and Stockholm Cancer Society ((#241282 to RL) and (#221383 to KV)). Professor Kantimay Das Gupta performed profilometry measurements on the SU-8 masters. The authors thank Senjuti Chakraborty, Afrah Aboo, Apoorva S. Raman, Rahul Shobhawat, Savita Kumari, Ramiz Raza, Kanak Joshi, Anushka Bairoliya and Pradip Shinde for technical help. The authors would also like to thank Professor Odd Terje Brustugun, Vestre Viken Hospital Trust, Drammen and Institute for Cancer Research, Oslo University Hospital, Oslo Norway and his team for providing the H1975/OR cell line used in some of the experiments. Some of the chemicals used in this work were kindly donated by M. P. Biomedicals through a MoU with WRCB, IIT Bombay. The authors acknowledge Professor Shamik Sen, Professor Swati Patankar, Prof. Prasenjit Bhaumik and Dr. Abdur Rub for their valuable suggestions on this work. The authors thank the IIT Bombay Nanofabrication Facility (IITBNF), the Centre for Sophisticated Instruments and Facilities (CSIF), and the Spinning Disc Confocal Microscope facility of IIT Bombay. The common instrument facility of the Department of Biosciences and Bioengineering, IIT Bombay, is acknowledged for the use of cell culture facility, ultracentrifuge, gel documentation system, multimodal plate reader and the dynamic light scattering measurement system. A part of Figure 1 was prepared using BioRender.

