## Supplementary Information for "LoAR-Exo and HiAR-Exo: One-step label-free isolation of extracellular vesicles using inertial microfluidic devices"

### HiAR-Exo: a high-aspect ratio inertial microfluidic device for one-step label-free isolation of extracellular vesicles from biological samples

Sourav Acharya[1], Moein Talebian Ghevari[2], Ashwin Anandhapadman[1], Shreya Karandikar[1], Haimanti Mukherjee[1], Fredrik Stridfeldt[3], Sayani Das[1], Petra Hååg[4], Nupur Agarwal[4], Rolf Lewensohn[4,5], Kristina Viktorsson[4], Sandip Kaledhonkar[1], Prakriti Tayalia[1], Apurba Dev[2] and Debjani Paul[1]

[1]Department of Biosciences and Bioengineering, Indian Institute of Technology Bombay, Mumbai, India

[2]Division of Solid-State electronics, Department of Electrical Engineering, Uppsala University, Uppsala, Sweden
[3]Department of Applied Physics, Kungliga Tekniska Högskolan Royal Institute of Technology, Stockholm, Sweden
[4]Dept of Oncology and Pathology, Karolinska Institutet, Stockholm, Sweden
[5]Theme Cancer, Patient area Head, Neck, Lung and Skin Cancer, Karolinska University Hospital, Stockholm, Sweden

#### 1. Theory of particle separation in spiral microchannels

##### 1A. Theory of particle separation in low-aspect ratio inertial microfluidics

Following the conditions required for focusing of particles in an inertial microfluidic system as given by Di Carlo [Di Carlo. 2009]. The flow-rate required to focus particles at an equilibrium position is given by

$$Q = \frac{2\pi\mu w^3 h}{3\rho L a^2 f_L}$$

Here, $\mu$ is the viscosity of the fluid, w is the width of the channel, $h$ is the height of the channel, $L$ is the length of the channel, $a$ is the particle diameter, $f_L$is the lift coefficient.

In the LoAR-Exo design, we have $w$ is 75 µm and $h$ is 66 µm (AR ~1). The fluid properties ($\rho$ and $\mu$) are considered as the fluid properties of water. L is 8 cm and $f_L$ is 0.05 [Di Carlo. 2009]. For focusing particles > 1 µm, we calculated the required flow rate for particles having a diameter of 2 µm and 3 µm. The required flow rates for focusing of 2 µm and 3 µm particles in their equilibrium positions are 212 µL/min and 91 µL/min. We consider a flow-rate of 150 µL/min as the flow-rate for focusing of particles > 1 µm. We then conducted fluid flow simulation at a flow-rate of 100 μL/min and a radius of curvature (R) of 5 mm to check the formation of stable dean vortices.

In addition to that, in a curved microchannel, the ratio of inertial lift force to dean drag force ($R_f$) should be > 0.04.

$$R_f = \frac{a^2 R}{w^3}$$

For particles having size 2 μm and 3 μm, the value of $R_f$ are 0.048 and 0.108 respectively. We can use LoAR-Exo for focusing and removal of particles >1 μm from the sample. Now, the above conditions are applicable for aspect ratios of 0.5 to 2 [Di Carlo. 2009.]. For focusing of particles in a high aspect ratio spiral microchannel (AR>2), we have used the theory developed by Cruz. et al. [Cruz et al. 2021].

#### 1B. Theory of particle separation in high-aspect ratio inertial microfluidics

Following the derivation by Cruz et al [Cruz et al. 2021] the lift force ($F_L$) and Dean drag force ($F_D$) acting on any particle in the inertial flow are given by the following equations.

$$F_L = \frac{f_L \rho U_m{}^2 a^4}{w^2} \quad (1)$$

$$F_D = 3\pi\mu a v \quad (2)$$

Here, $f_L$ is a constant that depends on the position of a particle along the channel width, and has a value between 1 and 2. $\rho$ is the density and $\mu$ is the viscosity of the fluid. a is the size of particle, w is the width of the channel, $U_m$ is the maximum fluid velocity and $v$ is the particle velocity relative to the fluid.

As both forces are dependent on particle size, we can achieve sequential inertial focusing of particles in the same cross-sectional plane according to their size in a high-aspect ratio channel. To focus particles of a specific size at a particular position and concentrate them, the lift force acting at the focus region must be in the same order as the Dean force at the focus region. In our first design iteration, we considered width and height of the channel as 50 μm and 100 μm respectively. We have simulated flow rates ranging from 100 μL/min to 500 μL/min. Stable dean vortices are attained only at 500 μL/min. The lift force and Dean drag force acting on a 1 μm particle are calculated for a flow rate of 100 μL/min. The maximum fluid velocity ($U_m$) and the order of magnitude of the particle's relative velocity ($v$) were determined from COMSOL simulation results. The value of $U_m$ for a flow rate of 100 μL/min is 0.66 m/s whereas the order of magnitude of $v$ in the secondary flow direction is $10^{-3}$ m/s.The fluid properties ($\rho$ and $\eta$) are considered as the fluid properties of water. The lift force is in the order of $10^{-14}$ N whereas the Dean drag force is in the order of $10^{-12}$ N. Evidently the lift force is not adequate to restrict 1 μm particles near the inner walls. This is confirmed by the particle simulation result where the device is incapable of focusing 1 μm particles.

The lift force is inversely proportional to the width of the channel. So in our second design iteration, we reduced the width of the channel to 20 μm to increase the lift force. We then calculated the lift force and Dean drag force acting on 1 μm particles. The value of $U_m$ for a flow rate of 100 μL/min is 3.22 m/s whereas the order of magnitude of $v$ in the secondary flow direction is $10^{-2}$ m/s. Both the forces are in the order of $10^{-11}$ N. It fulfills the necessary condition to focus particles near the inner wall of the channel.

We then optimized the radius of curvature (R) and the aspect ratio (AR) using COMSOL simulations, where different R values were tested for the formation of stable Dean vortices. The simulation results identified R = 5 mm as the optimal radius of curvature for a flow rate of 100 µL/min.

#### 2. Optimization of imaging conditions in NTA for small and large particles

We report our observations on avoiding some common pitfalls during NTA measurement. As NTA is an optical method, the scattered light from the particles in the solution is directly correlated with their size. Therefore, using one single setting for light intensity and the camera gain can misrepresent the size distribution in a heterogeneous sample containing both nano and micron-sized particles. For instance, a high intensity setting of the light and a high gain setting of the camera are required to characterize small particles as they have a weak scattering of light compared to large particles. However, using this same setting for the entire sample mixture would scatter too much light from the large particles, saturating the camera and hiding the small particles. We set both the brightness and the sensitivity values as 'low' for measuring the concentration of 1 µm particles, while keeping both brightness and sensitivity values as 'high' for measuring the concentration of 100 nm particles.

#### 3. COMSOL simulation for optimization of device design and flow parameters in HiAR-Exo

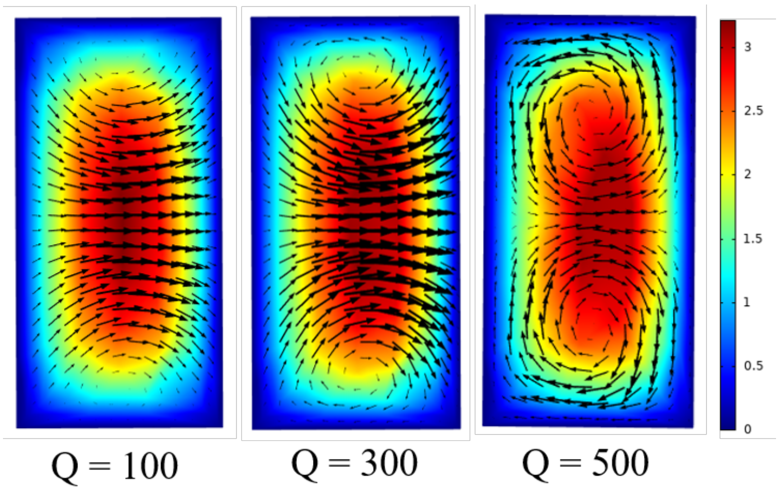


*Figure S1. COMSOL simulation showing the streamline distribution over the channel cross-section for a radius of curvature of 10 mm. Streamlines are shown for three flow rates of 100 µL/min, 300 µL/min and 500 µL/min. Stable counter-rotating vortices are seen only at a flow rate of 500 µL/min.*

**Figure S1** shows the streamlines from COMSOL simulation in the cross sectional view of HiAR-Exo for an R = 10 mm. Here it is observed that at Q = 100 µL/min and Q = 300 µL/min, there are no vortices formed inside the channel. Stable counter-rotating vortices are only formed at Q = 500 µL/min.

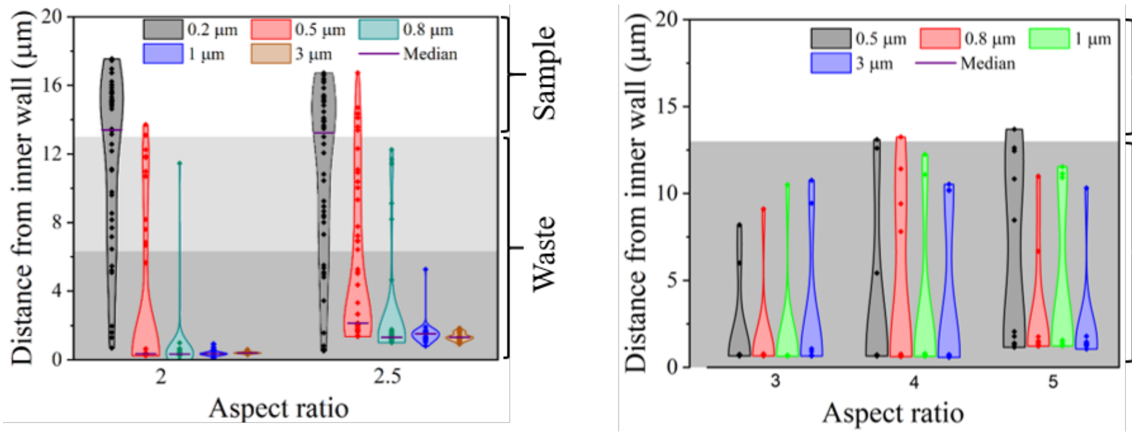


*Figure S2. Violin plot from COMSOL simulations showing the focusing distance from the inner wall of individual particles for an aspect ratio of 2, 2.5, 3, 4 and 5.*

We simulated the separation of polystyrene particles ($10^6$ particles/mL) with diameters 200 nm, 500 nm, 800 nm, 1 μm, and 3 μm using the particle tracing module of COMSOL. **Figure S2** (COMSOL simulation results) shows the violin plot of the distance of the individual particles from the inner wall for all the different values of the aspect ratio (AR) (AR = 2, 2.5, 3, 4, 5). At AR = 5, none of the particles focus at a precise location. Instead, they are randomly scattered across the cross section of the waste outlet. As particle size decreases, the lift force acting on them also becomes weaker. This increases the likelihood of 1 μm particles entering the sample outlet. A similar pattern is observed at AR = 4. When AR is reduced to 3, focusing improves for both 3 μm and 1 μm particles. These particles remain confined within the waste outlet area. However, 0.5 μm particles are still not observed in the sample outlet, which goes against the intended function of the device.

As shown in **Figure S2**, the COMSOL simulation results show that 0.8 μm, 1 μm and 3 μm particles are focused and collected from the waste outlet for aspect ratios of 2 and 2.5. For AR = 2, 5% of the 0.5 μm particles go to the sample outlet, while this number increases to ~ 16% for AR = 2.5. For both aspect ratios, 52% of 0.5 μm are directed to the sample outlet. Hence we fix the value of AR to 2.5 as we can collect both 0.2 μm and 0.5 μm in the sample outlet.

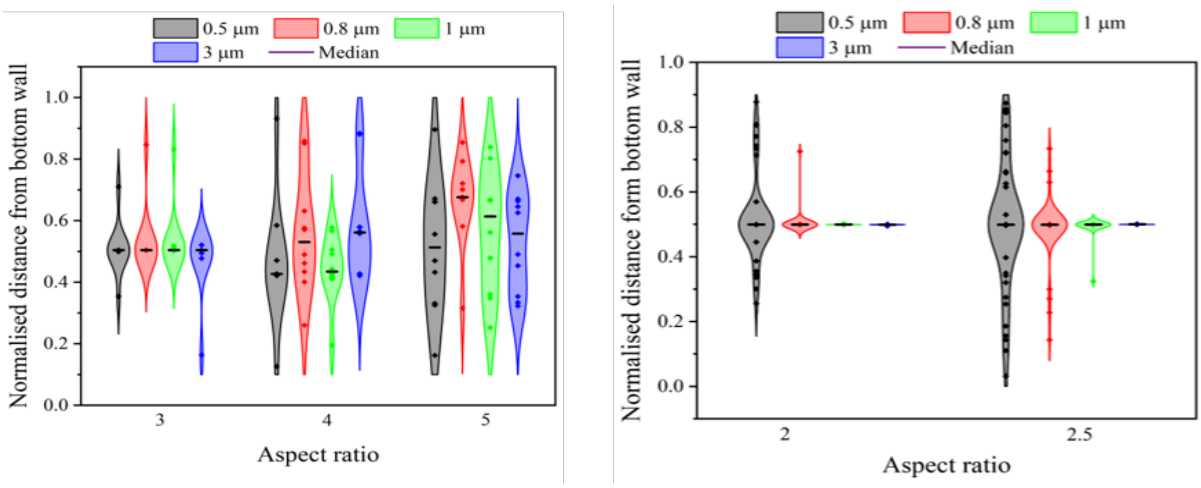


*Figure S3. COMSOL simulation results showing vertical focussing of particles. Normalized distances of 3 µm, 1 µm, 0.8 µm and 0.5 µm particles from the bottom surface of the channel for aspect ratios of 5, 4, 3, 2.5 and 2 are shown. Each particle position has been normalized by the respective channel height.*

We calculated the distance of particles from the bottom surface of the channel to evaluate the microchip's focusing efficiency in the vertical direction. **Figure S3** shows violin plots from COMSOL simulation showing the normalized vertical positions of the particles. The positions have been normalized by dividing with the appropriate channel height.

As the aspect ratio decreases from 5 to 3, the particle focusing becomes tighter. Precise vertical focusing of larger particles (3 µm and 1 µm) is observed at aspect ratios of 2.5 and 2. In contrast, 0.5 µm particles do not focus much in the vertical direction. These results suggest that the focusing behaviour in the vertical direction follows a trend similar to that in the horizontal direction. However, vertical focusing is less critical to the operation of HiAR-Exo, as particle separation primarily depends on their horizontal focusing positions when exiting the channel.

#### 4. Full images of the western blots

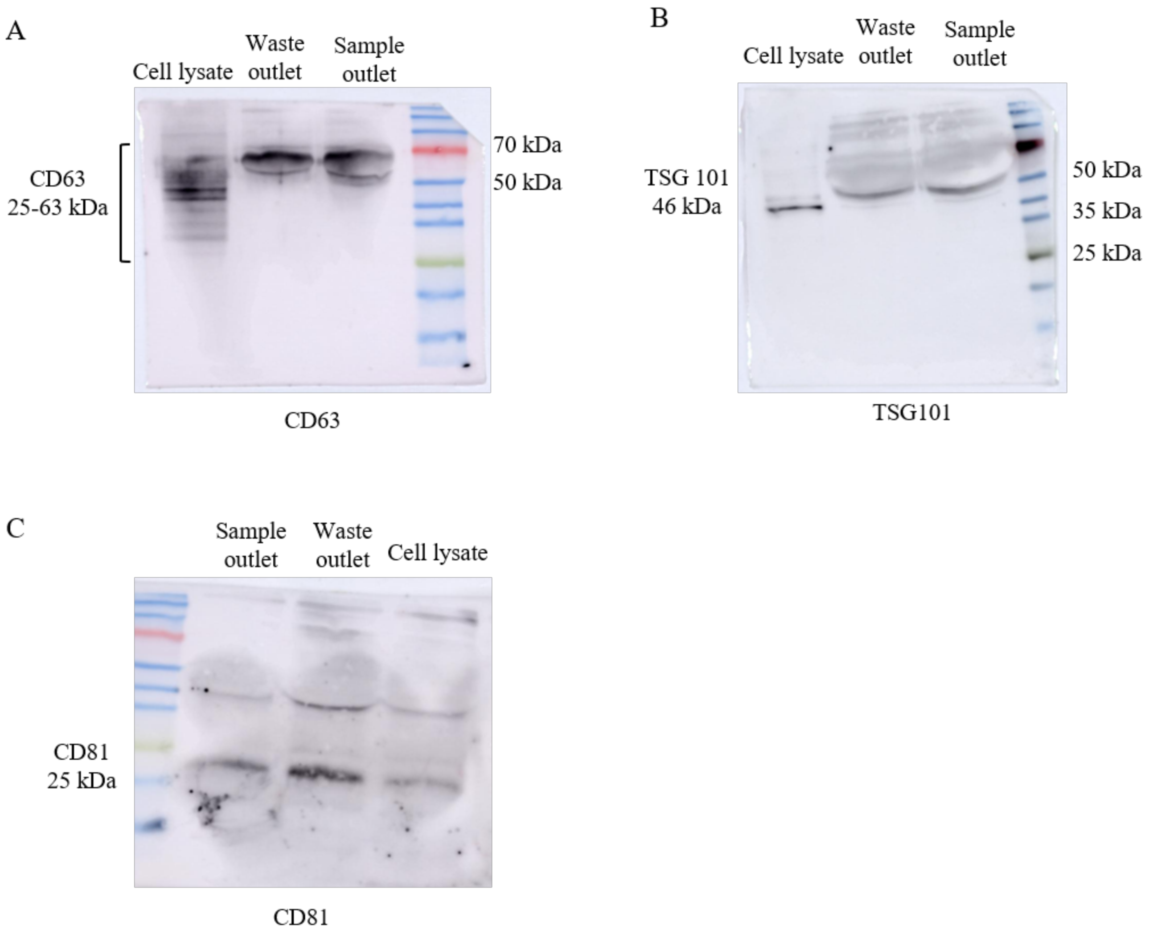


Figure S4. Full images of the western blots from CCM of MDA-MB-231. Blots for CD63, TSG 101 and CD81 are shown. EVs were  isolated using LoAR-Exo.

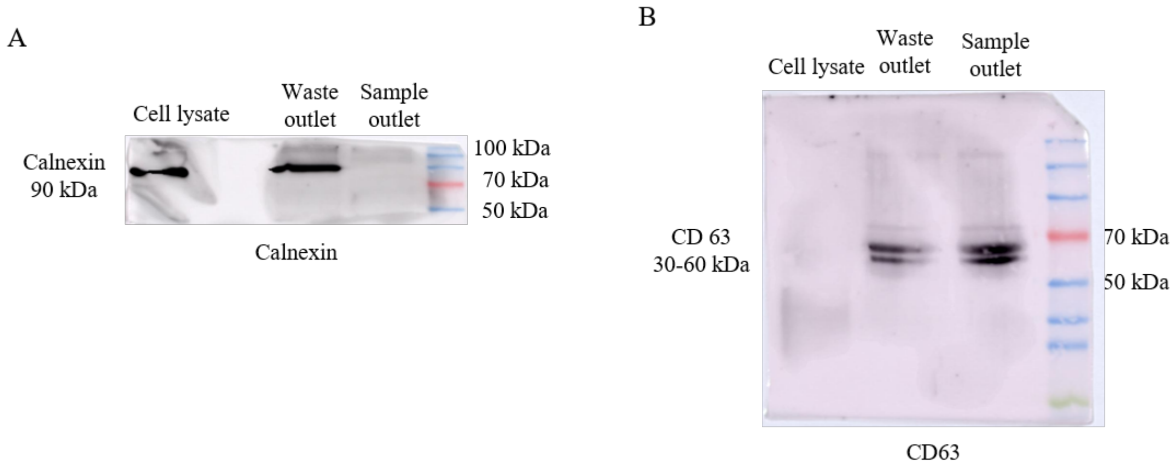


Figure S5. Full images of the western blots from cell suspension of MDA-MB-231 spiked in CCM using LoAR-Exo for isolation. Blots for calnexin and CD63 are shown.
